# Direct optical measurement of intra-molecular distances down to the Ångström scale

**DOI:** 10.1101/2023.07.07.548133

**Authors:** Steffen J. Sahl, Jessica Matthias, Kaushik Inamdar, Taukeer A. Khan, Michael Weber, Stefan Becker, Christian Griesinger, Johannes Broichhagen, Stefan W. Hell

**Affiliations:** Max Planck Institute for Multidisciplinary Sciences, Department of NanoBiophotonics, 37077 Göttingen, Germany; Max Planck Institute for Medical Research, Department of Optical Nanoscopy, 69120 Heidelberg, Germany; Max Planck Institute for Multidisciplinary Sciences, Research Group Structure and Dynamics of Mitochondria, 37077 Göttingen, Germany; University Medical Center Göttingen, Department of Neurology, 37075 Göttingen, Germany; Max Planck Institute for Multidisciplinary Sciences, Department of NMR-based Structural Biology, 37077 Göttingen, Germany; Max Planck Institute for Medical Research, Department of Chemical Biology, 69120 Heidelberg, Germany; Abberior Instruments America, Bethesda, MD 20814, USA; Leibniz-Forschungsinstitut für Molekulare Pharmakologie, 13125 Berlin, Germany

## Abstract

Optical investigations of nanometer distances between proteins, their subunits, or other biomolecules have been the exclusive prerogative of Förster Resonance Energy Transfer (FRET) microscopy for decades. Here we show that MINFLUX fluorescence nanoscopy measures intra-molecular distances down to 1 nm – and in planar projections down to 1 Angström – directly, linearly, and with Angström precision. Our method is validated by quantifying well-characterized 1-to-10 nm distances in polypeptides and proteins. Moreover, we visualize the orientations of immunoglobulin subunits and reveal specific configurations of a histidine kinase domain dimer. Our results open the door for examining proximities and interactions of macromolecules under physiological conditions.

---

Due to its minimal invasiveness, fluorescence microscopy has played a central role in the quest for understanding the myriad of functions of biomolecules. Understanding these functions, however, calls for the quantification of sizes, associations, and conformational changes of proteins and other biomolecules, which in turn requires precise measurements of inter- and intramolecular distances *r*=1-20 nm. While measuring such distances is clearly challenged by diffraction, superresolution fluorescence microscopy also struggles with this task. As a matter of fact, since the 1960s, macromolecular distances have been inferred indirectly, namely by the phenomenon called Förster Resonance Energy Transfer (FRET)^**1**^. Concretely, FRET deduces distances *r* from the probability with which an excited fluorophore transfers its excited state energy to a relaxed fluorophore featuring a longer excitation and emission wavelength that is located further away by *r*. As the only optical method to explore proximities <10 nm in both protein ensembles and on the individual protein level, FRET^**2-4**^ has become one of the most popular methods in the life sciences. Dubbed a ‘molecular ruler’, FRET has thus led to many insights both in cells and in vitro^5-7^.

On the other hand, FRET entails experimental provisions stemming from the rather indirect nature of the measurement^5^. Since the energy transfer occurs via an interaction of the two transition dipole moments of the fluorophores, the result varies inversely with the sixth power of the distance, 1⁄*r*^6^, rendering FRET a highly nonlinear ‘ruler’^6,7^. Moreover, FRET depends on the usually uncontrolled relative orientation of the two transition dipoles and on the polarizability of the molecules between them, altogether making reliable determinations of *r* challenging and every so often impossible. The situation is exacerbated by the fact that FRET measurements are actually limited to (at most) 2-8 nm, because the dipole-dipole interaction hardly modulates with *r* outside this range. All these factors can lead to considerable uncertainties, making FRET much better at providing ‘yes or no’ answers regarding co-localization, for example in large molecular ensembles and in cells, rather than at determining exact (intra)molecular distances.

In principle, nanometer-scale proximities between two spectrally shifted fluorophores with different excitation and/or emission wavelengths can be measured^8-10^, because the spectral shift allows for the separation of the fluorophores despite diffraction. Provided the spectral shift is sufficient, each fluorophore can be localized by calculating the centroid of its fluorescence diffraction spot rendered on a camera. Ironically, in the sought-after range of *r* ≲ 8 nm the spectral shift undesirably elicits FRET between the fluorophores, thus hampering their independent emission. Independence can be gained by ensuring that the fluorophores emit sequentially, through an on-off switching of their fluorescence, like in the superresolution^11^ method called PALM/STORM^12-14^. However, reversible on-off switching does not decouple the fluorophores entirely, and for distances < 10 nm this residual coupling^15^ unfortunately elevates bleaching^16^.

Improved emission independence can be gained by binding fluorophores transiently to the sites of interest, as in the method called PAINT^17^, which is highly effective when mediated by DNA hybridization^18^. Although sub-nanometer localization precisions can be obtained^19^, DNA-PAINT requires that the sites of interest can be decorated with and accessed by (labeled) DNA single strands. This labeling procedure is by far more restricting than directly attaching a fluorophore at the protein site of interest, and not all biomolecules can be labeled by DNA oligonucleotides. Moreover, fluorophore labels need to be constantly replenished by fresh labels in the surrounding solution, altogether making DNA-PAINT not viable in living cells.

Conceptually even more serious at this scale are the fundamental limitations of localizing a fluorophore by establishing the centroid of its fluorescence diffraction spot on a camera. First, centroid calculation critically relies on the detection of many fluorescence photons, since the localization precision *σ* scales with 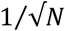, with *N* being the total number of detected photons in the camera spot. Thus, the need to wait for many emissions entails frequent excursions of the fluorophore to transient, non-emitting states, sample drift and bleaching. Second, unless the fluorophore emits isotropically in space, which is virtually impossible given that it is target-bound, the dipole nature of emission leads to systematic deviations of the centroid position from the actual position of the molecule^20-23^ that can exceed 10 nm. As the rotational freedom and the emission (an)isotropy are a priori unknown, these deviations mandate special caution regarding the accuracy of any method based on centroid determination, especially at molecular distances.

Here, we show that MINFLUX localization^**24**^ determines (intra)molecular distances linearly and with sub-nm precision at room temperature. In MINFLUX localization, the fluorophore position is established by relating, and possibly matching, the known position of a central zero of a doughnut-shaped excitation beam to the unknown position of the fluorophore (Fig. 1A). Thus, MINFLUX implies a minimization rather than a maximization of the number of required fluorescence photons. Because in typical MINFLUX localization the localization precision *σ* scales exponentially with the number of detected photons *e*^−*N*^, the issues of camera-based localization are overcome. In fact, MINFLUX typically requires about 100 times fewer detected photons to reach the camera localization precision^24,25^. By utilizing photoactivatable (caged) fluorophores, emission orthogonality can also be ensured such that one fluorophore emits while the other one remains fully inert to the excitation. Additionally, the localization of the fluorophores with a circularly polarized doughnut zero renders the localization virtually independent of the unknown orientation of their emission dipoles. The resulting reliable determination of each fluorophore’s coordinates has enabled us to measure molecular distances directly, that is, without mediation by a multi-factorial process such as FRET. Quantifying 1-20 nm distances has thus allowed us to visualize, for example, the end-to-end distance of a small 16 kDa protein or provide access to conformational details of a larger 150 kDa protein by direct positional measurements of the reporter fluorophores.

**Fig. 1.**
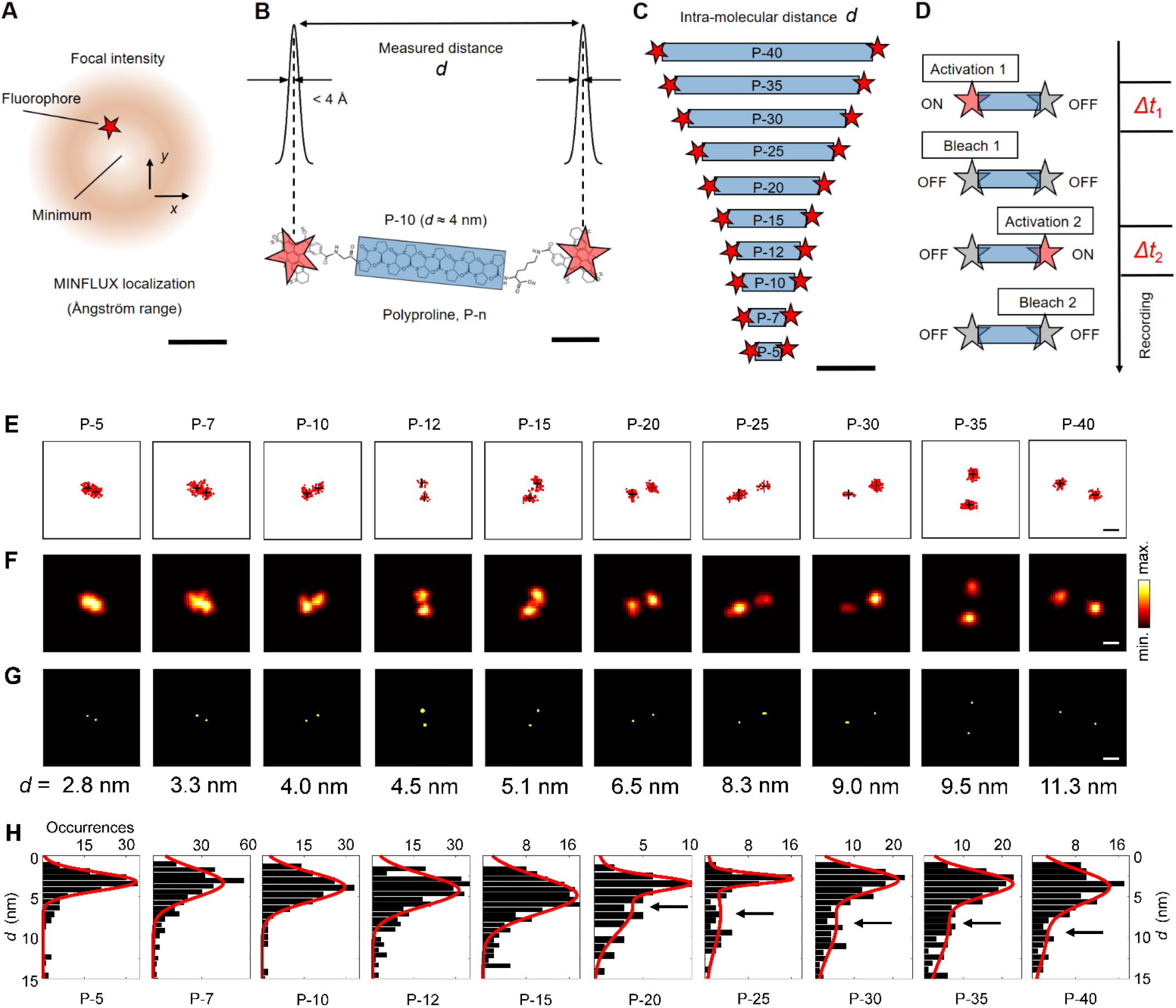
Ångström-precise intra-molecular distance measurements based on direct optical measurements of positions, in and beyond the FRET range. (**A**) MINFLUX approaches a fluorophore (red star, not to scale) with an intensity minimum (center of doughnut pattern) to achieve its precise localization. (**B**) Tunable distances provided by polyproline, a peptide comprised of relatively rigid helices of defined repeat number, terminal glycine and lysine residues, and identical photoactivatable fluorophores conjugated to both ends. The Angström-range MINFLUX localization precision enables measurements of 1-10 nm intra-molecular distances. (**C**) Prepared polyproline constructs: P-n, with n = 5,7,10,12,15,20,25,30,35,40, constituting a distance “ladder”. (**D**) Sequential photoactivation used for addressing the fluorophores individually and in molecular proximity. (**E**-**G**) MINFLUX reconstructions of polyproline end-to-end distances, displayed as (E) localization sets (red points) and the localized dye coordinates (black crosses), 2D histogram representation, and (G) inferred dye positions and their respective measurement precision. The yellow ellipses correspond to 3σ contours (or >3 Å for visibility). The extracted distance *d* is indicated below each example. (**H**) Distributions of measured distances *d* from P-5 to P-40, exhibiting >20% spread in the relative standard deviation that is attributed to considerable flexibilities in the proline polymer chains, terminal residues and fluorophores. For 20 and more residues, long distances can be discerned in a broad second peak (indicated by arrows), but most end-to-end distances observed on surface were found to be shorter (first peak). Scale bars: 200 nm (A), 1 nm (B), 3 nm (C), 5 nm (E-G).

## MINFLUX as a linear ruler in the <10 nm (intra)molecular range

To explore the potential of the chosen MINFLUX protocol (*Methods*, Fig. 1A), we initially turned to the polyproline helix secondary structure as a distance reference. The widely assumed rigidity of this polypeptide^26,27^, especially for low to intermediate numbers (<20) of repeats, allows to position fluorophores at defined distances. In fact, polyprolines served Stryer and Haugland to demonstrate^28^ the *1*⁄*r*^6^ FRET dependence in the 1960s, put forward by Förster in 1948. Polyprolines also allowed Schuler, Eaton and coworkers to demonstrate distance-dependent FRET at the single-molecule level^29^.

Conjugation of the photoactivatable dye DiMeO-ONB-SiR637 (ref. ^30^) to both ends of polyproline chains P-n with amino- and carboxyl-terminal glycine or lysine residues produced putatively linear unimolecular systems (Fig. 1B). The synthesis was controlled to yield peptides with negligible variation in the proline repeat number, meaning that the inter-fluorophore distance *d* was tunable from about 2 nm (for P-5) to >10 nm (for P-40), taking as an expectation the highly extended type II (PPII) helix lengths with all-trans peptide bonds (Fig. 1C). The peptides were unspecifically immobilized on the surface of a poly-L-lysine coated cover glass at high dilution and examined individually (see *Methods*).

To quantify the localization precision for individual fluorophores at the end of the polyprolines, we initially imaged samples of a separated fraction of mono-labeled P-15 polyproline. Fluorophores that exhibited noticeable positional drifts due to insufficient immobilization were excluded from analysis. Photoactivation was ensured by a regularly focused 405-nm beam that was co-aligned with the 640-nm doughnut-shaped excitation beam. Following activation of DiMeO-ONB-SiR637 by cleavage of the *ortho*-nitrobenzyl carbamates, the fluorophore was transferred from an inactive off-state to an active on-state with photostable red emission. Once an active fluorophore was identified, the MINFLUX localization algorithm ensured that the excitation doughnut rapidly zoomed in on the emitter^31,32^. The final two MINFLUX iterations^32^ were repeated until photobleaching, rendering extended sets of successive fluorophore position estimates (Fig. S1). From these raw localization data, the statistical localization precision can be inferred directly. For example, for only 100 photons in the final MINFLUX iteration, we obtained *σ*_raw_ = (2.2 ± 0.2) nm. Aggregation of groups of 5 and 10 consecutive localizations yielded datasets with a smaller spatial spread, namely *σ*_5 combined_ = (1.2 ± 0.2) nm and *σ*_10 combined_ = (0.9 ± 0.2) nm, respectively.

The mean position of the fluorophore, i.e. its time-averaged center of mass, is subject to an even lower uncertainty due to the combined information from many equivalent position determinations obtained for long on-times. In practice, obtaining on average >1000 localizations, the localization precision was estimated to be on the order of the microscope’s drift stabilization per measurement time interval (0.1-0.3 nm). Under the assumption of stationary fluorophore centers, the mean of all localizations can be assigned with even (sub-)Angström precision depending on numbers of localizations accrued (Fig. S1). Note that our stabilized MINFLUX system^32^ including beamline monitoring did not require further post-processing of localization data to correct for drifts.

Moving to polyprolines with two DiMeO-ONB-SiR637 dyes, we observed that, apart from a fraction of already activated fluorophores at the beginning of the measurement, the emitters were indeed activated and localized independently (Fig. 1D). Localization of a fluorophore was terminated by a photobleaching event. As the activation probability, controlled by the 405-nm laser power, was kept low, the activation of the second fluorophore frequently occurred seconds to minutes later, showing that the activation of the two fluorophores was indeed independent.

### Precise and accurate distance measurements in the FRET range

MINFLUX allowed to quantify the fluorophore-fluorophore distance (Fig. 1E-G) from P-40, with examples in the range *d ≈* 10-12 nm, over intermediate spacings (e.g. P-30; *d ≈* 8-9 nm) down to spacings deep within the FRET range (e.g. P-15; *d ≈* 5 nm); the shortest tested peptide was P-5 (*d ≲* 3 nm). The sub-nanometer precision of individual position measurements produced data that were easy to evaluate even at small *d*.

The distributions of measured distances (Fig. 1H) were generally broad, and especially so for the longer polyprolines. This broadening, interpreted to result from a considerable flexibility in the peptide chain, had also been observed in the single-molecule FRET data and was described in simulations^29^. For oligomers with *n ≥* 20, we observed rather broad peaks in the respective range expected for largely extended helices from ∼6 nm to ∼10 nm (highlighted by arrows in Fig. 1H). A large subpopulation of polyprolines, however, exhibited much shorter end-to-end distances contained in the respective first peak of the distribution. We speculate that these may arise in part due to pronounced bending and uncontrolled interactions with the surface, differing from the solution-phase environment of previous studies.

Up to 20 P, the average measured distances exhibited a close-to-linear scaling with the number of proline residues (Fig. 2A). Linear regression analysis in this range (P-5 to P-20) suggested an increment of Δ*d ≈* 0.21 nm per proline residue. While molecular dynamics simulations would explain moderate reductions of average end-to-end distance^29^ also for oligomers in this size range due to chain bending, the observed rise of – on average – 2.1 Å per residue is in disagreement with complete type II (PPII) conformation of the helices in their entirety. For such all-*trans* bonding in PPII conformation, which we expect to be generally favored under our experimental conditions, the helical pitch of 9.3 Å / turn and 3.0 residues / turn would result in a rise of 3.1 Å per residue. Instead, our data (Fig. 2A) point to heterogeneous compositions of *cis* and *trans* peptide-bond structural segments which in their combination lead to the average remarkably linear scaling observed. Clearly, subpopulations of *cis*-prolyl isomers interspersed in the chain reduced the average end-to-end distances of the chains^33,34^.

**Fig. 2.**
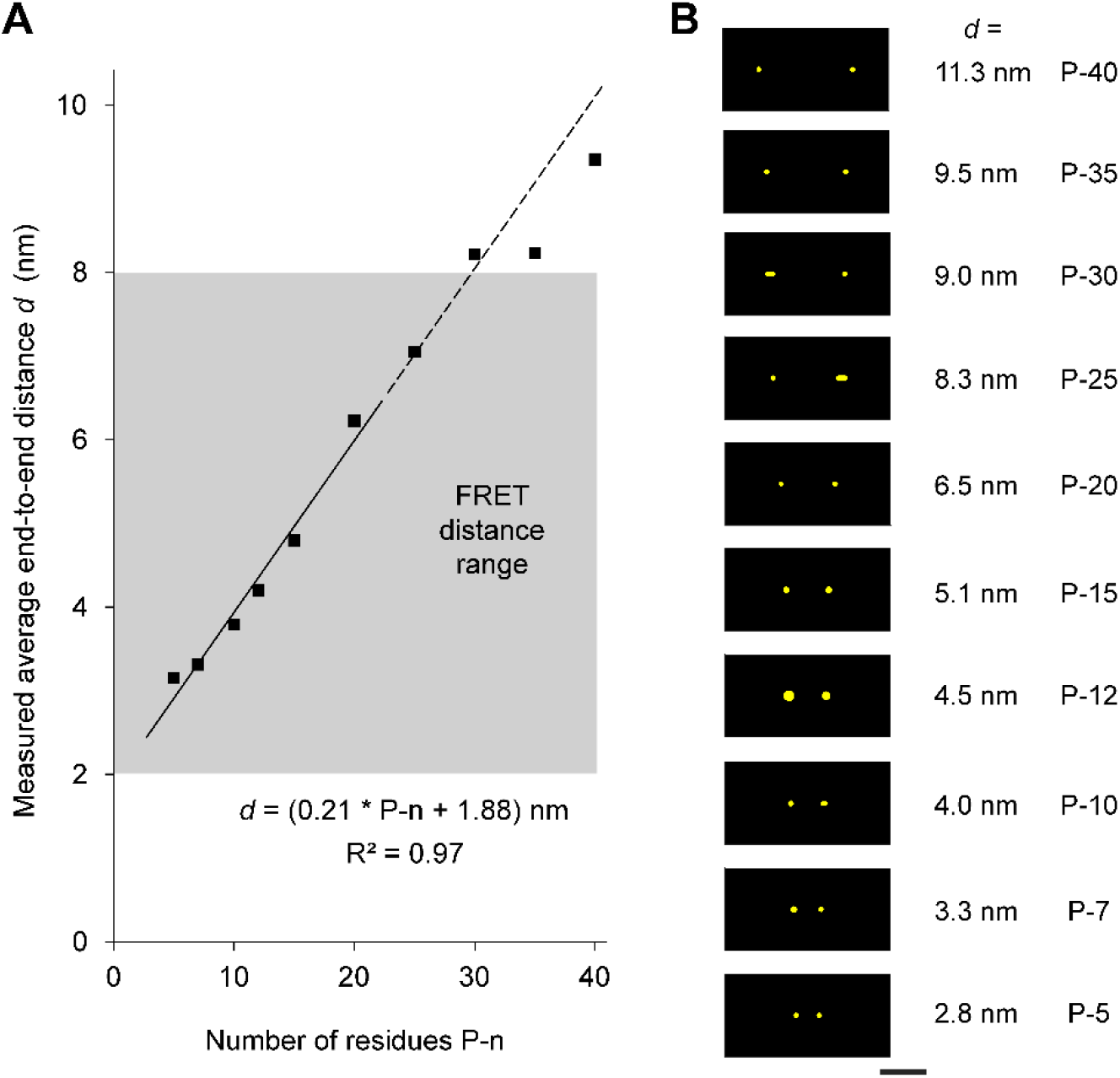
Intra-molecular MINFLUX imaging of polyproline end-to-end distances. (**A**) Measured average distances vs. number of proline residues P-n. For P-5 to P-15, the averages were extracted as the positions of the peaks of the distributions in Fig. 1H. For P-20 and longer, an indication of the average more extended distances is given from the positions of the second, broader peaks. A linear fit over the P-5 to P-20 range is shown, suggesting a rise of, on average, 2.1 Å per residue. This suggests heterogeneous compositions of *trans* and *cis* peptide bonding in the polyproline helix. The range of distances ∼2-8 nm has been traditionally addressed by (single-molecule)FRET methods (grey region). (**B**) Example data from Fig. 1 E-G, aligned by rotation to show the complete distance “ladder” (compare Fig. 1C). The ellipses represent 3σ contours of localization uncertainty (or >3 Å for visibility). Scale bar: 5 nm.

The constant offset (*y* intercept) of ∼1.9 nm may be attributed in part to the fluorophores’ displacement by the terminal glycine and lysine residues which act as linkers, and also to the fluorophores’ physical size contributing additional distance to the ‘centers’ of emission at both ends. In general, further contributions to distance variabilities may stem from non-horizontal attachments to the substrate and the 2D projection measurement. Altogether, a horizontal alignment of example data shows that the entire “ladder” of intra-molecular distances can be resolved and quantified (Fig. 2B vs. 1C) by MINFLUX.

### Distance measurements on a small protein

Building on this, we turned to examine C- and N-terminally labeled camelid nanobodies for end-to-end distance measurements. The core of this compact 16 kDa protein positions its C and N termini at ∼3.7 nm distance^35^ (Fig. 3A). The termini were labeled with a less hydrophobic photoactivatable dye^36^ by maleimide-coupling to the terminal cysteines. MINFLUX readily resolved this spacing, measuring (projected) distances in the range of 2.5 to 5 nm (Figs. 3B). The distribution of distances (Fig. 3C) peaks at ∼3.8 nm.

**Fig. 3.**
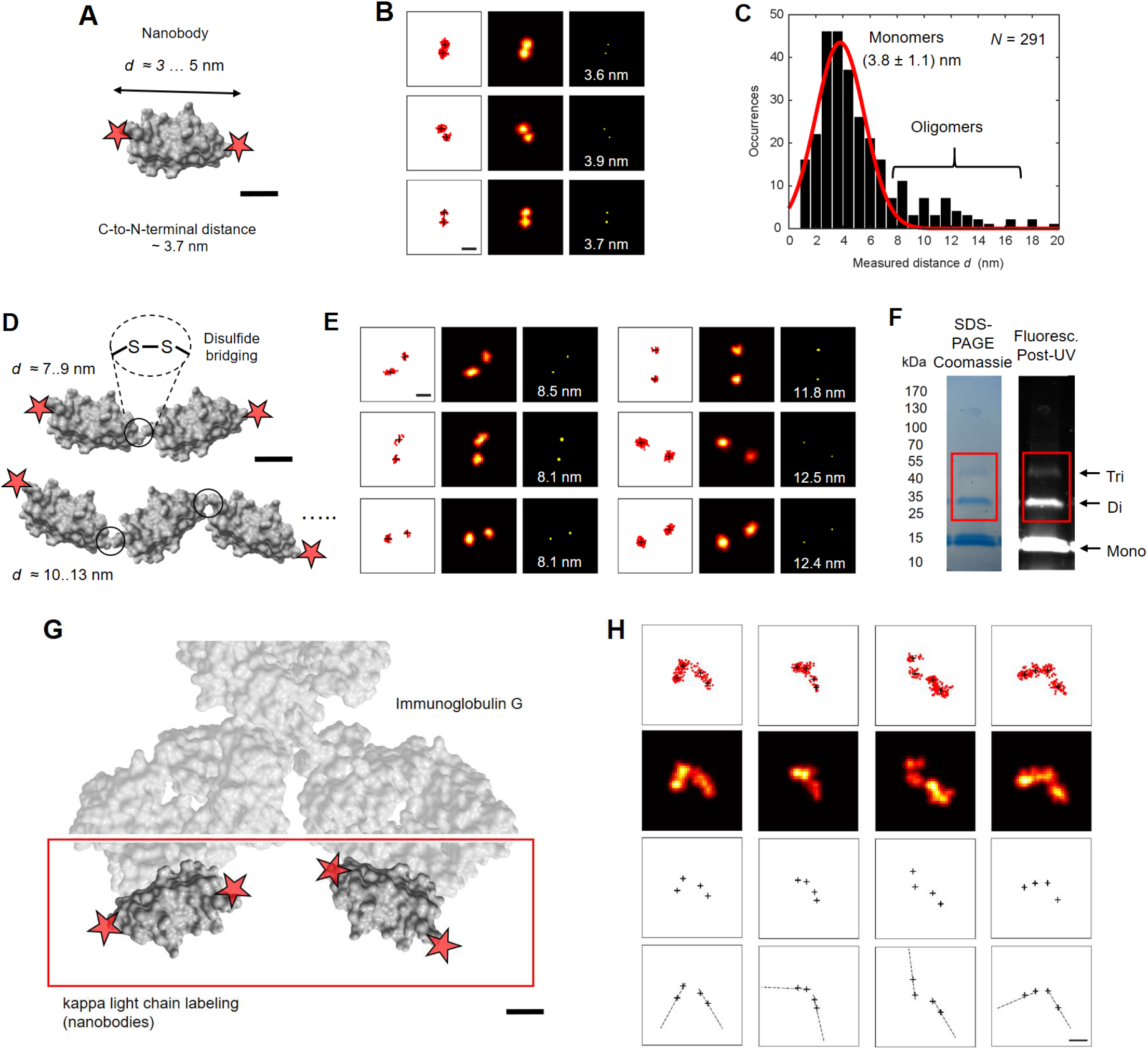
Intra-macromolecular distance measurements: sites on proteins and positioning of protein subunits. (**A**) Camelid nanobody (∼16 kDa) with N and C termini cysteine-labeled with a photoactivatable dye. The dimensions of the prolate-shaped nanobody core along the longer and shorter axis are ∼4 by ∼3 nm, and its N and C termini are 3.7 nm apart based on crystallographic data^25^. (**B**) MINFLUX example data. (**C**) Distribution of distances *d* obtained. Distance data in the long tail of the distribution suggest the existence of dimers and trimers. (**D**,**E**) Longer distances measured are attributed to oligomers (dimers and trimers). (**F**) Coomassie stain and fluorescence of a gradient gel of the solution as imaged, with dimer and trimer bands highlighted. (G) Immunoglobulin G molecule, with the kappa lights chains decorated by N-and C-terminal dye-labeled nanobodies, thus providing up to four positional marks on the antibody molecule. (H) Examples of different orientations of the two IgG “arms”. Scale bars: 1 nm (A,D,G), 5 nm (B,E,H).

Interestingly, the distribution is accompanied by a tail of longer spacings, extending out to ∼16 nm in rare instances. In particular, distances in the range of 7 to 9 nm, and 10 to 13 nm (examples in Fig. 3E) indicated the presence of nanobody dimers and trimers, whose end-to-end distances are expected to fall in this range. We hypothesized that dimers or trimers are formed by disulfide bridging (Fig. 3D,E) of the free cysteines during the maleimide labeling protocol. A gradient-gel SDS-PAGE analysis indeed revealed dimers and trimers both in Coomassie stain and in the fluorescence observed following UV photoactivation (Fig. 3F and Fig. S2). Owing to its high spatial resolution, MINFLUX obviously identified dimers and trimers directly by distance determinations. Note that the spacing of dimers and trimers falls outside the FRET range, highlighting the ability of MINFLUX to measure distances > 10 nm as well.

### Subunit structural arrangements of a large protein

Next, we extended the validity of our method to more reporter fluorophore sites. To this end, we labeled immunoglobulin G (IgG) with C- and N-terminally dye-labeled nanobodies having affinity for the IgG’s kappa light chain. This led to up to four fluorophores to be localized on the surface of the IgG. The fluorophores formed essentially two pairs aligned with either of the two rather flexible “arms” of the IgG (Fig. 3G). Due to incomplete labeling and/or sampling (Fig. S3A), not all fluorophore positions could always be extracted. Prolonged observation of individual dyes also indicated incomplete immobilization, as suggested in examples of relative displacements of one arm during acquisition (Fig. S3B). Yet, the antibodies for which all four dyes were registered clearly displayed the orientation of the two IgG arms (Fig. 3H).

### Dimeric protein domains in parallel and antiparallel configuration: down to Ångström distances

One of the exciting prospects of the newly gained resolution capability in the intra-molecular range is the direct imaging of distance distributions from an ensemble of macromolecules. Structure is not just one given rigid, non-flexible and non-varying arrangement, because molecules are flexible and this flexibility should become visible in the many snapshots acquired. The resulting dataset is expected to contain the mean as the dominant structure, which is also observed by the averaging pursued in crystallography or cryo-electron microscopy. However, as the imaging of individual molecules maps out their conformational space, structural subpopulations become apparent by MINFLUX.

To demonstrate this capability, we investigated a protein domain dimer, the cytosolic PAS domain (PASc) of the bacterial citrate sensor histidine kinase (CitA)^37^. Such a homodimeric system would be especially difficult to study by FRET, as the approach typically relies on two different dyes. Crystallography data of PASc reveal that this dimer can assume both anti-parallel and parallel configurations^38^ (Fig. 4A). A pronounced quaternary structure rearrangement in the form of an anti-parallel to parallel transition upon citrate binding is now thought to have a central role in transmitting and amplifying an initially small structural change as part of the transmembrane signaling process^38^.

**Fig. 4.**
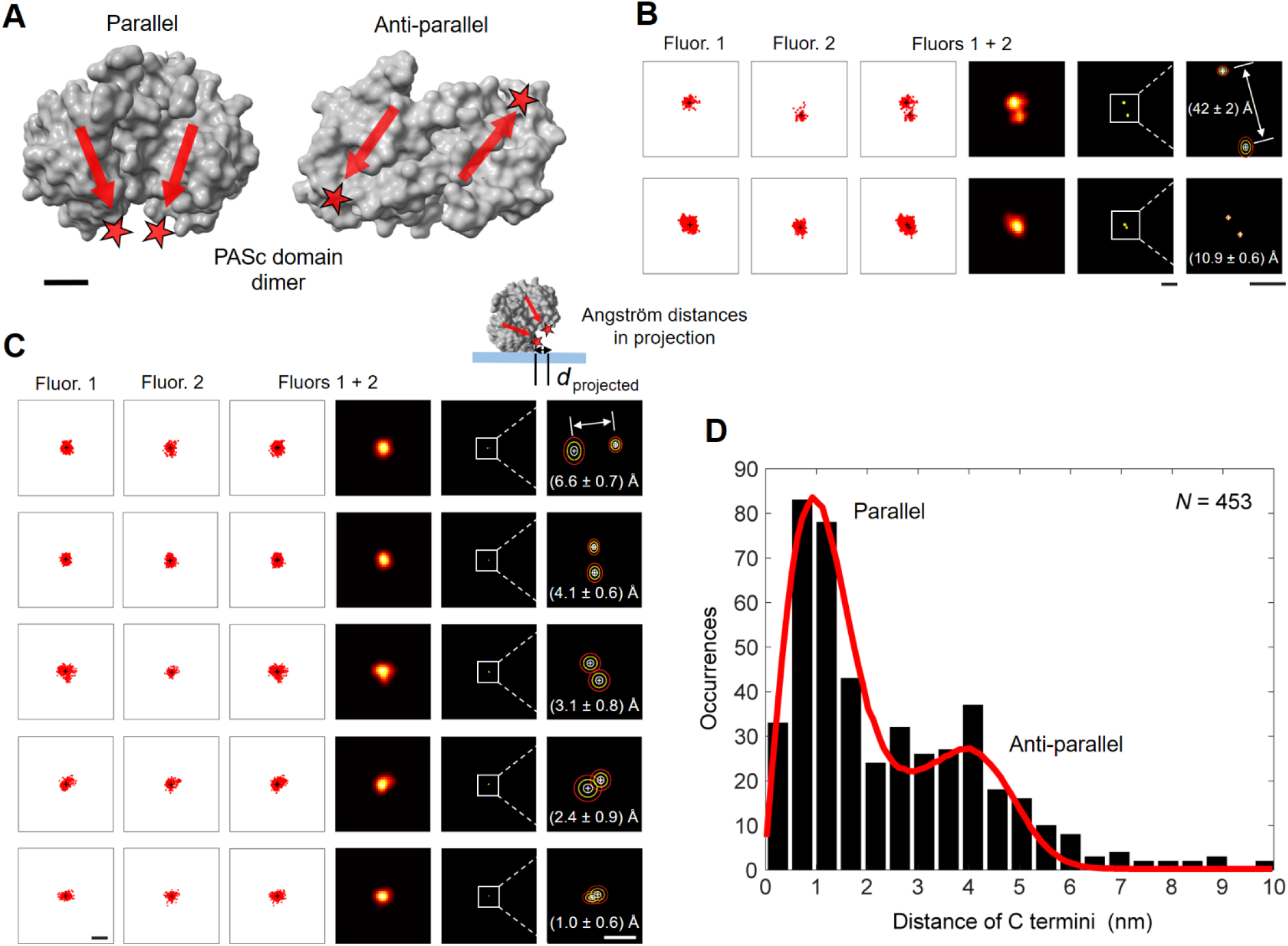
Conformational measurements at and below FRET-range distances, and Angström distance measurements in planar projections: Protein domain dimer in parallel and antiparallel configuration. (**A**) Crystallography data of the PASc domain of the bacterial citrate sensor histidine kinase (CitA) reveal that this dimer of protein domains exists in both anti-parallel and parallel configuration. The separation of the C termini, labeled with two photoactivatable dyes, differs between ∼1 nm for the parallel dimeric arrangement and ∼4 nm for the antiparallel dimer. (**B**) The set of distances observed include large numbers at ∼4 and ∼1 nm. (**C**) Projected instances lead to distances < 1 nm. The ellipses in the right enlarged views represent the 1σ (white), 2σ (yellow) and 3σ (red) contours of the position measurement uncertainty. (**D**) Complete distribution of measured (2D-projected) C-to-C terminal distance. Simulated projected distance distributions to capture the effects of the putatively random orientation of the dimers on the glass substrate can be found in Fig. S5. According to this simple modeling with the assumption of isotropic orientations (red curve, see *Materials and Methods*), the data indicate a mixture of about half parallel, half antiparallel. Scale bars: 1 nm (A), 5 nm and 2 nm (enlarged) (B), 5 nm and 5 Å (enlarged) (C).

The distance between the C-termini, labeled with photoactivatable dyes^36^, is expected to differ between ∼1 nm (∼10 Å) for the parallel dimeric arrangement and ∼4 nm for the antiparallel dimer. In our imaging experiments following sequential photoactivations (Fig. S4), distance data were observed ranging from shortest distances <1 nm, representing the parallel dimer at its full distance (or partially 2D-projected and thus reduced in length), to distances in the range of 4 nm, corresponding to the antiparallel arrangement (Fig. 4B). To appreciate the effects of the putatively random orientation of the dimers on the glass substrate, we simulated distributions of 2D-projected distances for species mixtures of various proportions (Fig. S5). The data clearly contain a population of tilted dimers, for which the axis between the two fluorophores has a perpendicular component to the plane, and for which distances between 0 and 1 nm were extracted with sub-Angström measurement precision (Fig. 4C).

The complete distribution (Fig. 4D) shows that about equal amounts of measured dimers were in the parallel and antiparallel state. MINFLUX thus quantified the relative abundance of the subpopulations with average C-terminal distances of ∼1 and 4 nm.

## Conclusion

MINFLUX fluorophore localization down to ∼0.1 nm precision enables accurate distance measurements down to the ∼1 nm physical extent of the fluorophores (Fig. 4C). For fluorophores positioned on macromolecules that lie at an angle and hence do not obstruct each other along the distance to be measured, we quantified distances even well below 1 nm. Resolving the end-to-end distance of small proteins and oligomers, these measurements provide optical access to the intra-macromolecular scale. As illustrated by distinguishing distances of 1-4 nm in parallel and antiparallel dimer configurations, MINFLUX also directly resolves subunits of larger proteins, as well as their relative orientation and conformation.

Our distance measurements further show that Ångström resolution can be *directly* obtained by observing just the two fluorophores, without the need for hundreds of copies of replenished fluorophores to bind to the two sites and define their coordinates over a long time.

MINFLUX enables linear distance measurements not only over the FRET range (2-8 nm) but also below 2-3 nm, where FRET usually falters. Also for distances >8 nm, MINFLUX measurements are fully viable (Fig. 3D,E). In fact, any distance from large, say 200 nm, to smallest, down to below 1 nm is directly measurable, because MINFLUX treats all distances equally.

With photoactivation as the on-off mechanism, we further observe that no “10-nm resolution barrier”^16^ was encountered, as for the STORM-type thiol-based blinking^39^, nor did the approach require cryogenic temperatures^40,41^. Unlike sample expansion^42^, which involves harsh chemical treatment, MINFLUX does not risk sample alterations at the sub-10-nm scale. The power of MINFLUX stems from the fact that it i) employs established procedures of fluorescence labeling, ii) uses the fluorescence photon budget about 100 times more effectively than conventional camera-based localization, while also avoiding iii) molecular-orientation effects and iv) FRET dipolar coupling. A currently remaining advantage of camera-based FRET imaging is the large field of view, offering a high degree of spatial parallelization, allowing for the examination of many molecules simultaneously. However, ongoing efforts for parallelization of MINFLUX should translate the virtues of MINFLUX to larger fields of view and living cells. Last but not least, our work shows that fluorescence microscopy is undergoing a seminal transition from a method that merely maps out molecular spatial distributions to one that will directly reveal their function with minimal invasiveness.

## Supporting information

Article

## Author contributions

S.J.S. initiated and led the experimental exploration of emitter co-localization (direct position measurements) in the FRET distance regime and below. S.J.S. and J.M. designed the imaging experiments. J.B. provided early input on the molecular model systems. S.J.S., K.I. and J.M. performed the imaging experiments. S.J.S. performed the data analysis, simulations and interpretation. T.A.K. provided the photoactivatable silicon rhodamine fluorophore. M.W. performed the gradient gel analysis and advised on photophysical properties of the activatable dye. S.B. and G.C. provided the GtCitA PASc domain dimers and structural comparisons by crystallography. S.W.H. conceived and developed the MINFLUX concept for ultra-precise, emission-photon-efficient localization, and provided critical feedback during the project. S.J.S. and S.W.H. wrote the manuscript, and all co-authors discussed the results and approved the final version of the manuscript.

## Corresponding authors

Correspondence to Steffen J. Sahl and Stefan W. Hell.

## Acknowledgement

We gratefully acknowledge excellent technical support at the MPI for Multidisciplinary Sciences (MPI-NAT) and the MPI for Medical Research (MPI-MR), and are grateful to S. Jakobs (MPI-NAT, University Medical Center Göttingen (UMG) and Fraunhofer Institute for Translational Medicine and Pharmacology) for generous access to a MINFLUX system (Abberior Instruments) funded by the German Research Foundation (DFG, grant no. INST 186/1303-1 to S. Jakobs). Jürgen Bienert, Jens Schimpfhauser and Jan Seikowski (all of the Facility for Synthetic Chemistry, MPI-NAT) coupled dyes to the proline polypeptides. S. Fabritz (MPI-MR) performed MS analysis. E. Rothermel (MPI-NAT) performed maleimide labeling of the nanobodies. F. Opazo (UMG) advised on nanobody labeling. K. Giller (MPI-NAT) produced the GtCitA PASc domain coupled with a dye. R. Schmidt (Abberior Instruments GmbH) helped with early experiments on the optical setup later reported in ref.^32^. D. Jans (UMG and MPI-NAT) provided further guidance and support with MINFLUX imaging. T.A. Hensel (MPI-NAT) gave helpful input on change-point detection analysis methods. V.N. Belov (MPI-NAT) advised on chemical aspects, and helped coordinate the dye coupling to the polypeptide rulers.

## References

1. Förster, T. Zwischenmolekulare Energiewanderung und Fluoreszenz. Annalen der Physik 437, 55–75 (1948).

2. Ha, T. et al. Probing the interaction between two single molecules: fluorescence resonance energy transfer between a single donor and a single acceptor. Proceedings of the National Academy of Sciences 93, 6264–6268 (1996).

3. Schütz, G.J., Trabesinger, W. & Schmidt, T. Direct Observation of Ligand Colocalization on Individual Receptor Molecules. Biophysical Journal 74, 2223–2226 (1998).

4. Zhuang, X. et al. A Single-Molecule Study of RNA Catalysis and Folding. Science 288, 2048–2051 (2000).

5. Roy, R., Hohng, S. & Ha, T. A practical guide to single-molecule FRET. Nature Methods 5, 507–516 (2008).

6. Stryer, L. Fluorescence Energy Transfer as a Spectroscopic Ruler. Annual Review of Biochemistry 47, 819–846 (1978).

7. Algar, W.R., Hildebrandt, N., Vogel, S.S. & Medintz, I.L. FRET as a biomolecular research tool — understanding its potential while avoiding pitfalls. Nature Methods 16, 815–829 (2019).

8. Churchman, L.S., Ökten, Z., Rock, R.S., Dawson, J.F. & Spudich, J.A. Single molecule high-resolution colocalization of Cy3 and Cy5 attached to macromolecules measures intramolecular distances through time. Proceedings of the National Academy of Sciences 102, 1419–1423 (2005).

9. Pertsinidis, A., Zhang, Y. & Chu, S. Subnanometre single-molecule localization, registration and distance measurements. Nature 466, 647–651 (2010).

10. Niekamp, S. et al. Nanometer-accuracy distance measurements between fluorophores at the single-molecule level. Proceedings of the National Academy of Sciences 116, 4275–4284 (2019).

11. Hell, S.W. Far-Field Optical Nanoscopy. Science 316, 1153–1158 (2007).

12. Betzig, E. et al. Imaging Intracellular Fluorescent Proteins at Nanometer Resolution. Science 313, 1642–1645 (2006).

13. Rust, M.J., Bates, M. & Zhuang, X. Sub-diffraction-limit imaging by stochastic optical reconstruction microscopy (STORM). Nat Meth 3, 793–796 (2006).

14. Hess, S.T., Girirajan, T.P.K. & Mason, M.D. Ultra-High Resolution Imaging by Fluorescence Photoactivation Localization Microscopy. Biophysical Journal 91, 4258–4272 (2006).

15. Ha, T. & Tinnefeld, P. Photophysics of Fluorescent Probes for Single-Molecule Biophysics and Super-Resolution Imaging. Annual Review of Physical Chemistry 63, 595–617 (2012).

16. Helmerich, D.A. et al. Photoswitching fingerprint analysis bypasses the 10-nm resolution barrier. Nature Methods 19, 986–994 (2022).

17. Sharonov, A. & Hochstrasser, R.M. Wide-field subdiffraction imaging by accumulated binding of diffusing probes. Proceedings of the National Academy of Sciences 103, 18911–18916 (2006).

18. Dai, M., Jungmann, R. & Yin, P. Optical imaging of individual biomolecules in densely packed clusters. Nat Nano 11, 798–807 (2016).

19. Reinhardt, S.C.M. et al. Ångström-resolution fluorescence microscopy. Nature 617, 711–716 (2023).

20. Enderlein, J., Toprak, E. & Selvin, P.R. Polarization effect on position accuracy of fluorophore localization. Optics Express 14, 8111–8120 (2006).

21. Stallinga, S. & Rieger, B. Accuracy of the Gaussian Point Spread Function model in 2D localization microscopy. Optics Express 18, 24461–24476 (2010).

22. Engelhardt, J. et al. Molecular Orientation Affects Localization Accuracy in Superresolution Far-Field Fluorescence Microscopy. Nano Letters 11, 209–213 (2011).

23. Lew, M.D., Backlund, M.P. & Moerner, W.E. Rotational Mobility of Single Molecules Affects Localization Accuracy in Super-Resolution Fluorescence Microscopy. Nano Letters 13, 3967–3972 (2013).

24. Balzarotti, F. et al. Nanometer resolution imaging and tracking of fluorescent molecules with minimal photon fluxes. Science 355, 606–612 (2017).

25. Eilers, Y., Ta, H., Gwosch, K.C., Balzarotti, F. & Hell, S.W. MINFLUX monitors rapid molecular jumps with superior spatiotemporal resolution. Proceedings of the National Academy of Sciences 115, 6117–6122 (2018).

26. Cowan, P.M. & McGavin, S. Structure of Poly-L-Proline. Nature 176, 501–503 (1955).

27. Adzhubei, A.A., Sternberg, M.J.E. & Makarov, A.A. Polyproline-II Helix in Proteins: Structure and Function. Journal of Molecular Biology 425, 2100–2132 (2013).

28. Stryer, L. & Haugland, R.P. Energy transfer: a spectroscopic ruler. Proceedings of the National Academy of Sciences 58, 719–726 (1967).

29. Schuler, B., Lipman, E.A., Steinbach, P.J., Kumke, M. & Eaton, W.A. Polyproline and the “spectroscopic ruler” revisited with single-molecule fluorescence. Proceedings of the National Academy of Sciences 102, 2754–2759 (2005).

30. Weber, M. et al. Photoactivatable Fluorophore for Stimulated Emission Depletion (STED) Microscopy and Bioconjugation Technique for Hydrophobic Labels. Chemistry – A European Journal 27, 451–458 (2021).

31. Gwosch, K.C. et al. MINFLUX nanoscopy delivers 3D multicolor nanometer resolution in cells. Nat. Methods 17, 217–224 (2020).

32. Schmidt, R. et al. MINFLUX nanometer-scale 3D imaging and microsecond-range tracking on a common fluorescence microscope. Nat. Commun. 12, 1478 (2021).

33. Doose, S., Neuweiler, H., Barsch, H. & Sauer, M. Probing polyproline structure and dynamics by photoinduced electron transfer provides evidence for deviations from a regular polyproline type II helix. Proceedings of the National Academy of Sciences 104, 17400–17405 (2007).

34. Best, R.B. et al. Effect of flexibility and cis residues in single-molecule FRET studies of polyproline. Proceedings of the National Academy of Sciences 104, 18964–18969 (2007).

35. Götzke, H. et al. The ALFA-tag is a highly versatile tool for nanobody-based bioscience applications. Nature Communications 10, 4403 (2019).

36. Kolmakov, K. et al. Masked red-emitting carbopyronine dyes with photosensitive 2-diazo-1-indanone caging group. Photochemical & Photobiological Sciences 11, 522–532 (2012).

37. Salvi, M. et al. Sensory domain contraction in histidine kinase CitA triggers transmembrane signaling in the membrane-bound sensor. Proceedings of the National Academy of Sciences 114, 3115–3120 (2017).

38. Zhang, X.C. et al. Mechanism of sensor kinase CitA transmembrane signaling. bioRxiv, 2023.02.06.527302 (2023).

39. Dempsey, G.T. et al. Photoswitching Mechanism of Cyanine Dyes. Journal of the American Chemical Society 131, 18192–18193 (2009).

40. Weisenburger, S. et al. Cryogenic optical localization provides 3D protein structure data with Angstrom resolution. Nature Methods 14, 141–144 (2017).

41. Dahlberg, P.D. et al. Identification of PAmKate as a Red Photoactivatable Fluorescent Protein for Cryogenic Super-Resolution Imaging. Journal of the American Chemical Society 140, 12310–12313 (2018).

42. Shaib, A.H. et al. Expansion microscopy at one nanometer resolution. bioRxiv, 2022.08.03.502284 (2022).

