## Supplementary material for "Direct optical measurement of intra-molecular distances down to the Ångström scale": Article

### Contents

**Materials and Methods**

**Fig. S1 |** Single-molecule MINFLUX localization precision in the Ängström range.

**Fig. S2 |** Full SDS-PAGE gel of nanobody monomer and oligomer populations.

**Fig. S3 |** Immunoglobulin G: Incomplete sampling of reporter fluorophore sites, effects of imperfect immobilization.

**Fig. S4 |** Sequential photoactivation and fluorescence emission of proximal dyes.

**Fig. S5 |** Simulations of projected distance distributions for mixture of species with 11 Å and 45 Å dye-dye distances.

**Supplementary References**

### Materials and Methods

*Polypeptides and proteins.* Polyprolines P-n (n = 5,7,10,12,15,20,25,30,35,40), with glycine and lysine residues at the N- and C- terminus, were synthesized at high purity by the Thermo Scientific custom peptide synthesis service (Life Technologies GmbH, Darmstadt, Germany), and conjugated in-house to DiMeO-ONB-SiR637 at both ends. The camelid nanobody (single-domain antibody) anti-ALFA with N-/C-terminal cysteines was purchased unlabeled from NanoTag Biotechnologies GmbH, Göttingen, Germany. The immunoglobulin G (IgG) monoclonal light-chain specific mouse anti-rabbit antibody was purchased from Jackson ImmunoResearch, Cambridge, UK (cat. number 211-002-171). The camelid nanobody FluoTag-X2 anti Mouse KLC (clone 1A23) was also purchased unlabeled from NanoTag Biotechnologies GmbH, Göttingen, Germany. The wild-type GtCitA PASc domains, modified by site-directed mutagenesis to carry a C-terminal cysteine residue for dye conjugation, were created as described previously[1,2]. They were stored in a buffer containing 20 mM trisodium phosphate, pH 6.5, 50 mM sodium chloride, 0.5 mM EDTA, 0.5 mM Pefabloc and 0.02% sodium azide. The camelid nanobodies and PASc domains were labeled with the maleimide derivative of Abberior CAGE 635, using standard protocols. Labeled peptides and proteins were stored as aliquots at -80° C before thawing on ice and diluting for sample preparation and imaging. For the immunoglobulin visualization, mixtures with the dye-labeled FluoTag-X2 anti Mouse KLC nanobody in 5-fold molar excess were incubated for 2 hours at room temperature before imaging the solution contents that included a population of the nanobody-bound immunoglobulin.

*Sample preparation.* Aqueous solutions (with ~20-30% dimethylsulfoxide (DMSO) for polyprolines) of the respective peptide or protein were serially diluted in PBS (nanobodies, IgG), GtCitA buffer (PASc) or trifluoroethanol (polyprolines) to find appropriate concentrations for surface densities of $\lesssim 0.01\ /\ \mu m^2$ to enable separate imaging. For polyproline imaging, self-assembled flow channels (coverslip glued to microscope slide with double sided tape (Scotch®, 3M France)) were treated with 150 nm gold colloids (BBI Solutions, SKU EM.GC150/4) (10 min), rinsed with $ddH_2O$ and incubated with the polyproline solution for 10 min. After rinsing with trifluoroethanol, the channels were sealed with epoxy glue (Epoxy Quick Set, UHU). For nanobody, IgG and PASc imaging, standard microscopy cover glasses (18 mm diameter, No. 1.5H, Marienfeld, 0117580) were functionalized with poly-L-lysine solution (10 min) (0.1 % (w/v), Sigma P8920), rinsed with PBS, treated with 150 nm gold colloids (BBI Solutions, SKU EM.GC150/4) (10 min) and rinsed again with PBS. Subsequently, the protein solution was pipetted onto the cover glass and incubated for 10 min. Following a rinse with PBS, the resulting samples were mounted on a microscope slide with a small cavity (Marienfeld, 1320002) to hold trifluoroethanol and sealed with 2K silicone (eco-sil, 1300 9100, Picodent).

*Gradient gel.* Nanobodies labeled with Abberior CAGE 635 were analyzed using reducing and non-reducing SDS-PAGE gel. Before loading, the samples were heated for 2 min to 65 °C in reducing (50 mM Tris HCl pH 6.8, 100 mM DTT, 2% SDS, 0.01% bromophenol blue and 10% glycerol) and non-reducing (50 mM Tris HCl pH 6.8, 2% SDS, 0.003% Coomassie Brilliant Blue R-250 and 5% glycerol) buffer. Samples were loaded (1:2:3:4 relative protein mass) onto a 4-15 % gradient gel (Bio-Rad, 4561084). After electrophoresis in standard electrophoresis buffer (10x: 0.25M Tris HCl pH 8.3, 1.925M glycine and 1% SDS), the gel was stained with Roti Blue (Carl Roth, A152.1) over night, activated using 365 nm UV light and scanned (full gel shown in Fig. S2).

*MINFLUX nanoscopy.* Image data were recorded on an Abberior Instruments MINFLUX nanoscope[3] using the standard 2D acquisition protocol, with settings of 10% (<0.5 µW average power at the sample) for the activation laser (405 nm) and 5% (~25 µW) for the excitation laser (640 nm) applied as part of the MINFLUX localization routine. Typically three or more gold colloidal particles positioned around an at least $5\times5\ \mu m^2$ field of view served as fiduciary markers for correction of the live-feedback sample and beam stabilizations. Once adjusted, the monitored acquisition was observed to be long-term stable for hours, including overnight acquisition.

*Data analysis and display.* A MATLAB® script using the *dbscan* routine identified candidate clusters containing at least 250 localizations, which were subsequently analyzed individually. The presented quantitative analysis excluded clusters for which the localization sets assigned to individual fluorophores indicated imperfect immobilization on the glass substrate, which were clearly identifiable by a smear-out over time, an unusually wide localization scatter, or a combination of the two. The stabilized MINFLUX system[3] does not require further post-processing of localization data. Any apparent motion is due to actual fluorophore proper motion, with the microscope coordinate reference frame always remaining entirely fixed.

For distance determinations, individual aggregated (10×) localization sets assigned to one fluorophore were required to exhibit a spread < 2 * $\sigma_{10\ \mathrm{combined}}$ (compare Fig. S1) to be included. The tightly confined localization clouds typically observed for immobile peptides and proteins easily fell within this criterion. To ensure robust measurements, we also required the less sampled position coordinate to be derived from at least 100 raw localization events, which meant that the precision reached ~3 Å for the "worse" localization in the pair. The numbers of successive localizations contributing to either position measurement were typically (much) larger.

At the spatial precisions of our experiments, *kmeans* clustering-based assignment of the localizations to two identifiable groups followed by calculation of the respective means was an effective distance estimator for distances ≳ 2.5 nm. In practice, either group corresponded to localizations with a given trace id (*tid*) parameter, indicating a separately identified new fluorophore, but the procedure can include additional numbers of localizations in either group with additional *tid*s (e.g. due to signal intermittency). The *tid* parameter is assigned anew by the microscope to each newly detected fluorophore[3]. After the successful termination of a localization event (typically as a result of terminal photobleaching) or after abording an unsuccessful localization attempt, a new *tid* is assigned to the subsequent event.

Distances that were initially estimated in this way to fall below 2.5 nm, i.e. with (substantially) overlapping localization scatter at 10× localization aggregates, were re-evaluated by the more precise procedure of grouping the localizations by the *tid*. Identification of more than two reporter fluorophore sites (here: the four sites on immunoglobulin) was possible by spatial clustering, *tid*-based cluster assignment, or a suitable combination.

A subset of the acquired fluorophore-pair localization clusters were contained within just one *tid*, and this could be ascertained both for cases of clearly visible displacements (such as $d$ ≳ 2.5 nm and well above), but also for close spacings as low as ≲1 nm. For those cases, irrespective of distance, a selection criterion could be established based on an abrupt change in emission properties that appears to differentiate the first from the second fluorophore, putatively due to minute local environment differences felt by the dye. Based on the collection of 10× aggregated localizations, the time series defined by ($t_{i+1} - t_i$) of the timepoints of localizations $i$ exhibited a pronounced discontinuity in its gradient, which was readily extractable by a changepoint detection on the diff($t_{i+1} - t_i$) timeseries. This served to identify clusters of one *tid* that were in fact two fluorophores, and the identified index allowed to assign the two localization groups and determine the displacement $d$ between their means.

Throughout the manuscript, unless otherwise indicated, localization data are presented as 10-fold aggregates (red points), along with black crosses (+) to show the mean of localizations attributed to a given reporter fluorophore. Isolated localizations remote to the assigned localization clouds were not displayed for clarity of visualization. Additionally, smoothed 2D histograms with a red-hot lookup table show the localization density (pixel width: 7.5 Å). In the 2D histograms, a square-root transform on the pixel value was applied to partially compensate localization number differences for the two fluorophore sites in display. The polyproline and nanobody distance measurement distributions exclude extreme projection examples of apparent distances $d$ < 1.25 nm. To appreciate the Ångström-level of the localizations, an alternative rendering represents the fluorophores' coordinates as an ellipse, showing outer contours of 3$\sigma$ in the localization measurement.

*Simulations.* Distributions of 2D-projected distances of assumed mixtures of two species of different length $d_1$ and $d_2$ (compare Fig. 4D) were computed as sets of a total of 1E6 distances, divided into the respective $d_1$ / $d_2$ fractions and evaluated as (normalized) histograms. Following individual randomized positioning of the two endpoints separated by $d_1$ or $d_2$ in the 2D plane, the projection according to $d_{\text{project}}$ = | ($d$ * cos($\theta$) | with a randomized projection angle $\theta$ = rand[0..1] * $\pi$ / 2 was calculated, i.e. assuming a uniformly isotropic orientation. The resulting positions to compute the distance were further subjected to a Gaussian localization spread with a standard deviation of $\sigma_x = \sigma_y$ = 5 Å, i.e. significantly worse than the localization precision of our experiments, but allowing for some of the putative conformational freedom of the dye labels.

*Structures.* For illustration purposes, renderings of the following structures were included to convey the dimensions of the respective molecules: Nanobody (Fig. 3), PDB code 6I2G; Immunoglobulin (Fig. 3), PDB code 1IGY; PASc domain (Fig. 4): PDB code 5FQ1.

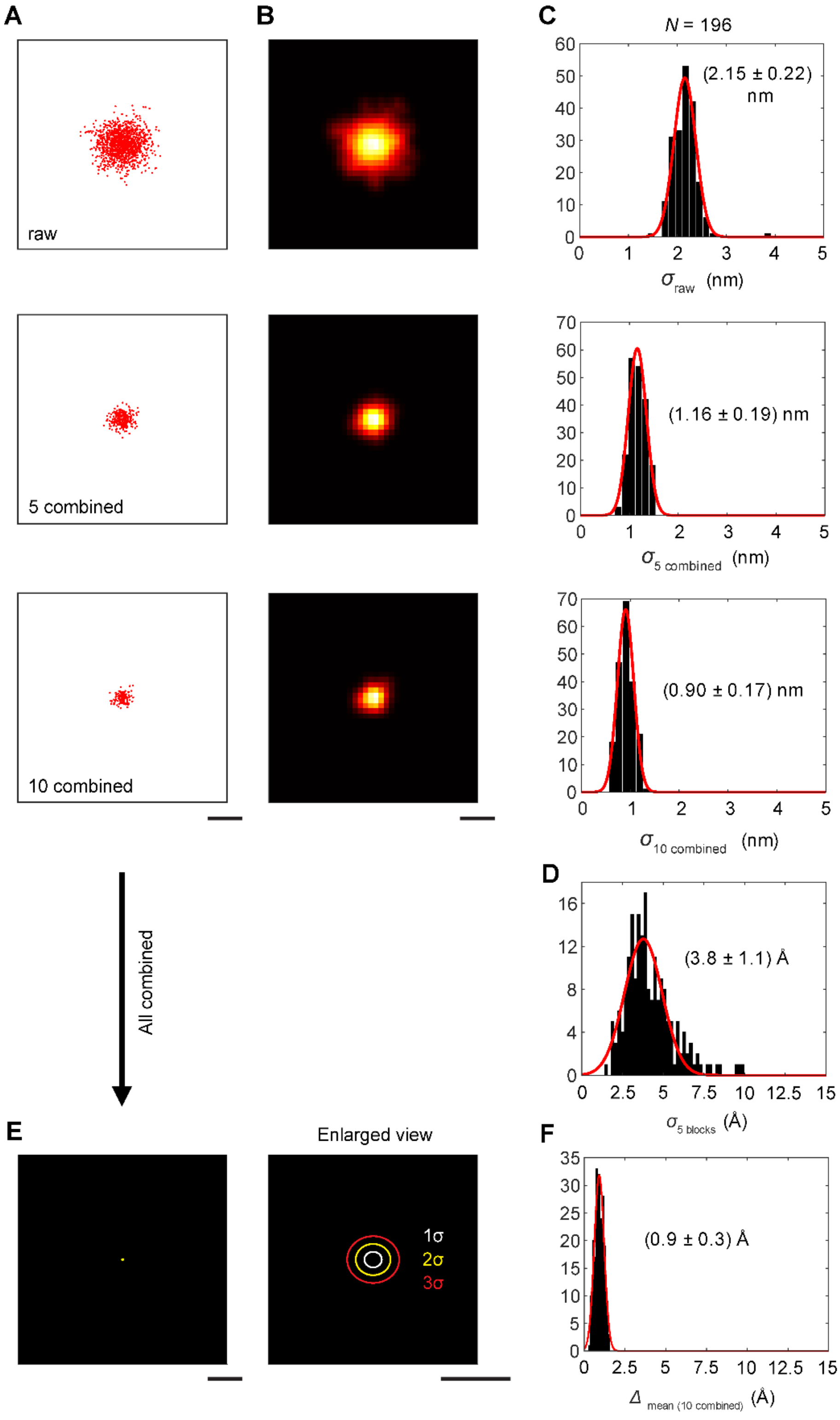


**Fig. S1 | Single-molecule MINFLUX localization precision in the Angström range.** Repetitive localization of single dyes from a mono-labeled polyproline sample (i.e. attached to a glycine/lysine linker under realistic intra-molecular measurement conditions). (**A**) MINFLUX localizations (raw, 5 and 10 sequential localizations combined). (**B**) Histogram representation to convey density of localizations. (**C**) Distributions of experimentally determined standard deviations. (**D**) Histogram of standard deviations of sets formed by combining localizations into 5 groups, yielding measured values of precision. (**E**) Under the assumption of stationary center of mass (c.o.m.), all localizations combined into a position estimate are subject to an uncertainty given by the error in the mean. The ellipses in the right enlarged views represent the 1σ (white), 2σ (yellow) and 3σ (red) contours of the position measurement uncertainty. (**F**) Histogram of calculated position uncertainty under assumption of stationary c.o.m. Scale bars: 5 nm (A,B,E), 5 Å (enlarged view in E).

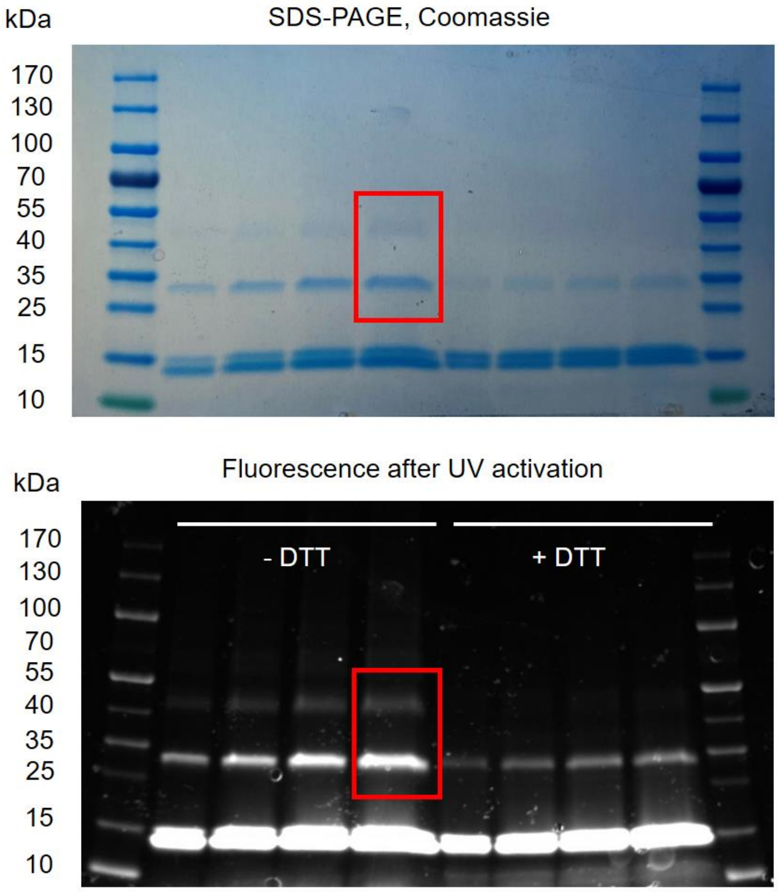


**Fig. S2 | Full SDS-PAGE gel of nanobody monomer and oligomer populations.**

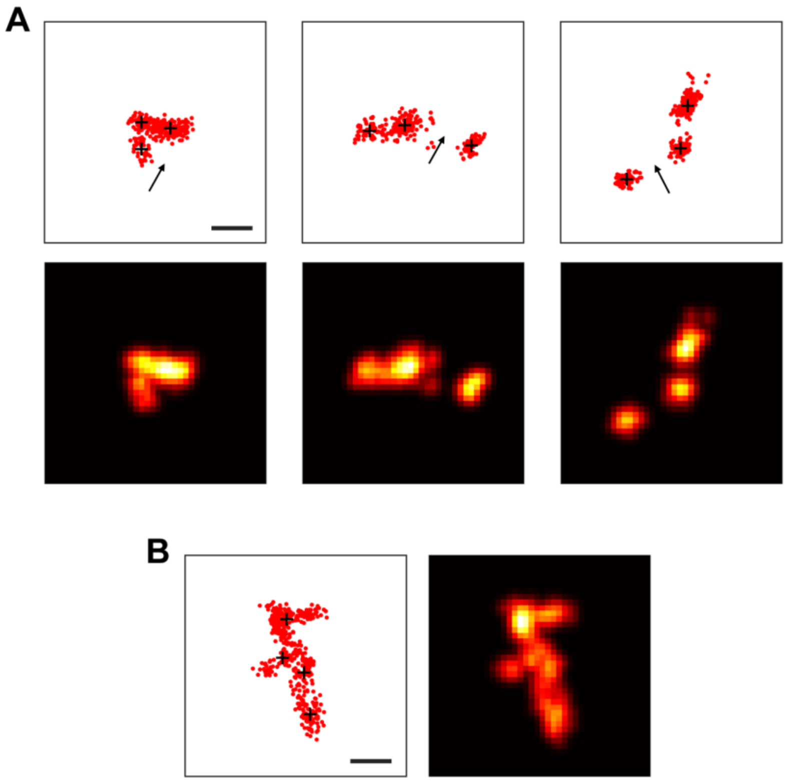


**Fig. S3 | Immunoglobulin G: Incomplete sampling of reporter fluorophore sites, and effects of imperfect immobilization.** (**A**) Examples of incompletely sampled IgG arms. The putative fourth position is indicated by an arrow in each case. (**B**) Example of motion of the IgG molecule during acquisition. Scale bars: 5 nm.

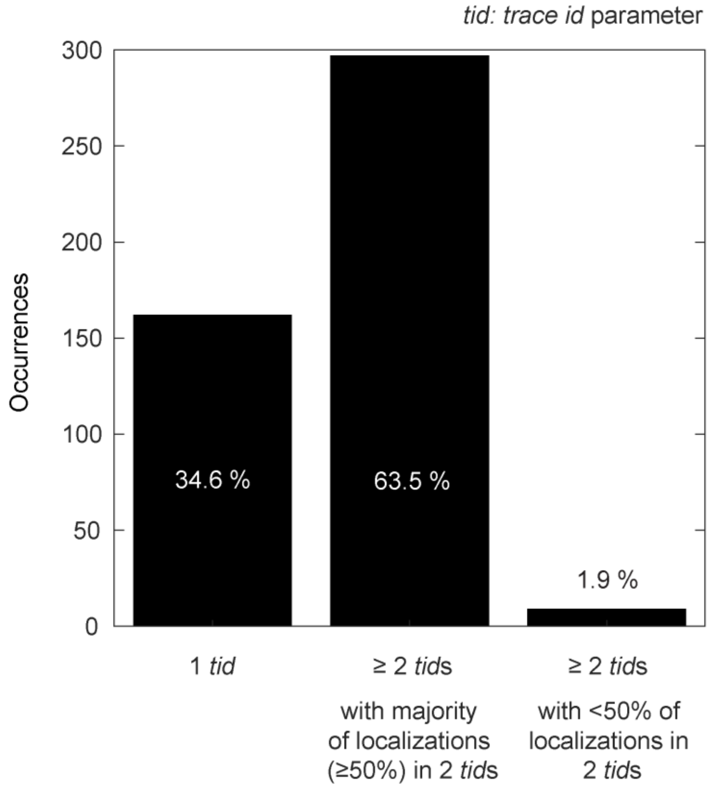


**Fig. S4 | Sequential photoactivation and fluorescence emission of proximal dyes.** Under conditions of low activation probability (low 405-nm power illumination), independent activation of the photoactivatable dyes leads to clearly identifiable time periods of emissions from either dye. The obtained localizations from a dye pair typically were comprised of two main groups assigned to two *trace ids* (*tid*s)[3], corresponding to the two dyes. Additional, fewer localizations carried additional *tid*s due to signal intermittency. For unambiguous assignment of fluorophores at small distance, the two main groups were used to yield the two sets of coordinates for the distance determination. A subset of localization clusters comprising localizations with only a single *tid* were inferred to subsume sequential emissions from the two dyes that could be confidently separated and assigned (see *Materials and Methods*). A negligible number of pairs featured larger numbers of *tid*s, in which case the assignment of the two most frequent *tid*s in the set, together a minority of localizations, still allowed a distance estimate, but at the cost of reduced spatial uncertainty due to lower numbers of combined localizations. Here, *tid* distribution data is presented for PASc domain dimers labeled at their C-termini (compare data in Fig. 4D).

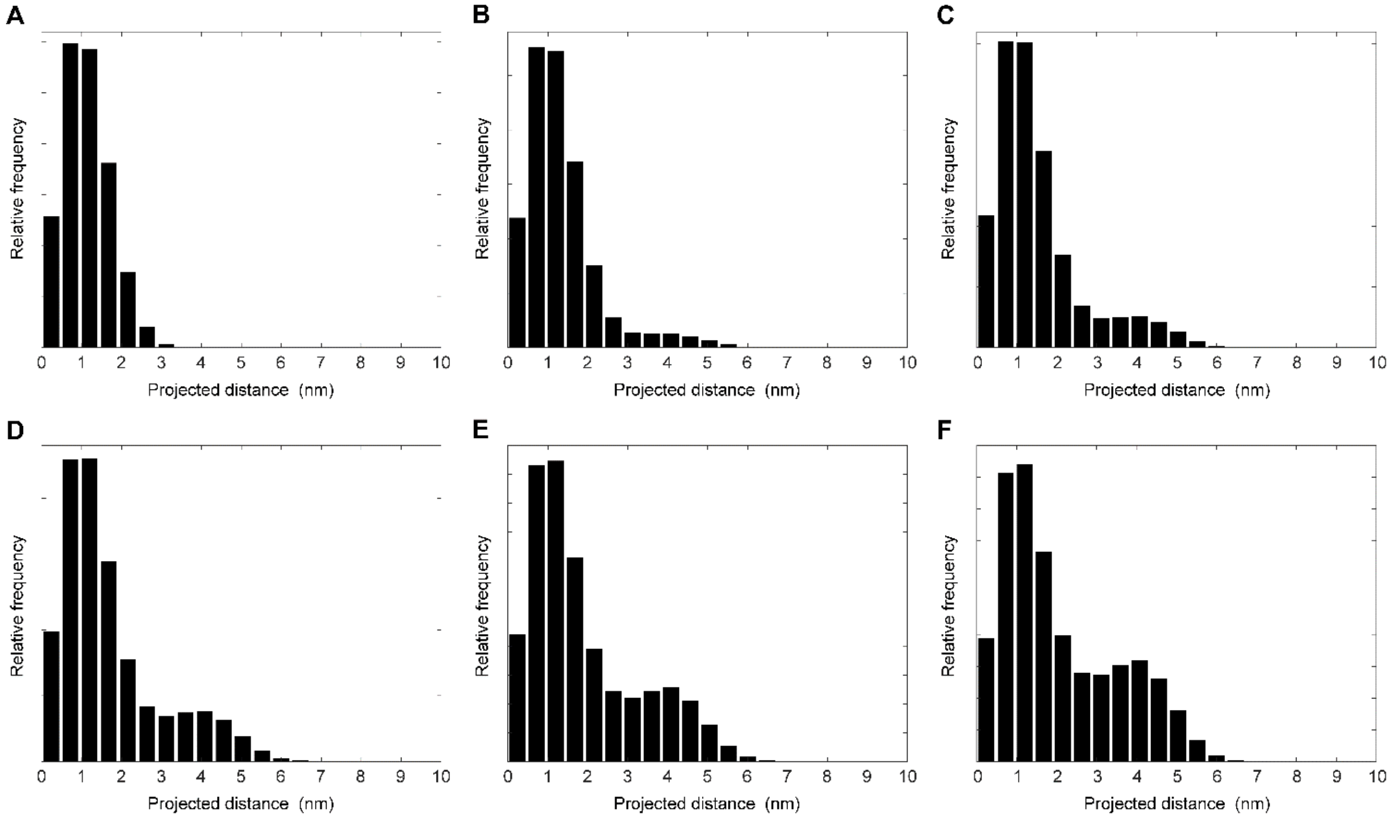


**Fig. S5 | Simulations of projected distance distributions for mixture of species with 11 Å and 45 Å dye-dye distances.** Projected distance distributions, under the assumption of isotropic orientation on the glass substrate, for (**A**) 100% of 11Å; (**B**) 90% of 11Å, 10% of 45Å; (**C**) 80% of 11Å, 20% of 45Å; (**D**) 70% of 11Å, 30% of 45Å; (**E**) 60% of 11Å, 40% of 45Å; and (**F**) 50% of 11Å, 50% of 45Å. See text and *Materials and Methods*.

### Supplementary References


1. Weisenburger, S. et al. Cryogenic optical localization provides 3D protein structure data with Angstrom resolution. *Nature Methods* **14**, 141-144 (2017).
2. Zhang, X.C. et al. Mechanism of sensor kinase CitA transmembrane signaling. *bioRxiv*, 2023.02.06.527302 (2023).
3. Schmidt, R. et al. MINFLUX nanometer-scale 3D imaging and microsecond-range tracking on a common fluorescence microscope. *Nat. Commun.* **12**, 1478 (2021).
